# Preventing photomorbidity in long-term multi-color fluorescence imaging of *S. cerevisiae* and *S. pombe*

**DOI:** 10.1101/180018

**Authors:** Gregor W. Schmidt, Andreas P. Cuny, Fabian Rudolf

## Abstract

Time-lapse imaging using multiple fluorescent reporters is an essential tool to study molecular processes in single cells. However, exposure to even moderate doses of visible excitation light can disturb cellular physiology and alter the quantitative behavior of the cells under study. Here, we set out to develop guidelines to avoid the confounding effects of excitation light in multi-color long-term imaging. We use widefield fluorescence microscopy to measure the effect of the administered excitation light on growth (here called photomorbidity) in yeast. We find that photomorbidity is determined by the cumulative light dose at each wavelength, but independent of the way excitation light is applied. Importantly, photomorbidity possesses a threshold light dose below which no effect is detectable (NOEL, no-observed-effect level). We found, that the suitability of fluorescent proteins for live-cell imaging at the respective excitation light NOEL is equally determined by the cellular autofluorescence and the fluorescent protein brightness. Last, we show that photomorbidity of multiple wavelengths is additive and imaging conditions absent of photomorbidity can be predicted. Our findings enable researchers to find imaging conditions with minimal impact on physiology and can provide a framework for how to approach photomorbidity in other organisms.

## 1 Introduction

Fluorescent live-cell imaging aims at using sufficient excitation light to obtain a detectable signal without influencing cell physiology [1,2]. Beside the fluorescent species of interest, a variety of endogenous and exogenous molecules are excited or absorb the excitation light [3]. Excited molecules can cause physiological disturbances due to chemical reactions or heating of the specimen. The nature of the involved molecules dictates what kind of damage arises. Mutagenic damage is often caused upon interaction of UV-B and -C light (λ <330 nm) with DNA and subsequent generation of cyclobutane pyrimidine dimers [4]. The stochastic nature of this damage can lead to effects at extremely low light doses and no threshold dose exists below which no effect is observed. Microscopists avoid exposing samples to short wavelengths by using dedicated UV blocking filters when illuminating with arc lamps, or by utilizing alternative light sources which do not emit UV light [5]. However, cellular damage including a different form of mutagenic damage can be caused by UV-A and visible light (>330 nm) through indirect mechanisms involving reactive oxygen species (ROS) [6,7]. ROS can be generated through the excitation of endogenous (e.g. metabolites) or exogenous (e.g. media components) light absorbing molecules (collectively termed photosensitizers) [3]. ROS can form adducts with nucleotides, proteins or other metabolites altering their function [3,7]. Cells are competent in clearing a limited amount of light-induced ROS damage as ROS are a bi-product of regular cell metabolism. Therefore, a threshold dose for light-induced damage exists, below which no effect on cell physiology is detectable [8].

How can the threshold light dose be determined? Initial attempts measured terminal phenotypes such as mitotic arrest in a tobacco cell line [9] or morphological changes and cell death in mammalian cells [10]. However, terminal phenotypes are not suitable to detect subtle photodestructive effects, which are expected to occur above a threshold light dose. Quantitative traits can be used to detect sublethal light-induced damage, for example, by measuring the nuclear localization frequency of the general stress-activated transcription factor Msn2 in *S. cerevisiae* [11,12]. In this case, a threshold light dose was found below which Msn2 localization frequency remained unchanged [12]. Disadvantages of this assay include its specificity to the *Saccharomyces sensu stricto*, and that it is unclear if the assay is equally sensitive to light of different wavelengths. Additionally, the assay relies on the measurement of a fluorescent signal which is related to the administered light dose and can be distorted by photobleaching. Alternatively, the growth rate (GR) in cell numbers provides a well-established, sensitive, quantitative and generic measure of the cellular well-being. For example, changes in GR have been used as a quantitative read-out in chemogenetic profiling [13] and systematic mapping of genetic interaction networks [14]. Also, it was previously proposed to measure the extent of light-induced damage in live-cell imaging by measuring changes in the GR of *S. cerevisiae* [15] or green monkey kidney cells [2]. Changes in growth, measured as the timing of cell division during early embryogenesis, were indeed shown to allow for quantifying subtle, non-toxic effects induced by fluorescence excitation light in *C. elegans* [8].

Here, we set out to analyse the confounding effects of visible light in multi-color long-term fluorescence imaging using the model organisms *S. cerevisiae* and *S. pombe*. We aimed to measure “photomorbidity”, defined as the decrease in growth rate upon light exposure, as a quantitative readout for sub-lethal effects of excitation light. We showed that photomorbidity can be described using a classical dose-effect relationship with a characteristic effective dose (ED50) and a no-observed effect level (NOEL) below which confounding effects are not detectable. We found that both measures depend on the cumulative light dose, and within practical boundaries are independent of the imaging interval, light intensity and bandwidth. To compare the suitability of different imaging channels and fluorescent proteins for live-cell imaging, we determined the signal-to-noise ratio (SNR) obtained with fluorescent fusion proteins at the NOEL of their specific excitation wavelengths. Our results demonstrate how photomorbidity and cellular autofluorescence, in addition to fluorescent protein brightness, can influence the performance of fluorescent proteins in live-cell imaging. We found that the combined photomorbidity of several wavelengths in multi-color imaging was additive and could be avoided by limiting light doses to the NOEL. Our findings highlight how particular combinations of fluorescent reporters enable artifact free multi-color time-lapse imaging. Guidelines on how to determine the NOEL light doses as well as an extended discussion of the optimization of time-lapse experiments can be found in the Supplementary Material.

## 2 Results

### Measuring photomorbidity in time-lapse microscopy

Decreased GR of cells is a well established indicator to measure adverse effects. We implemented a cell number based GR assay using a microfluidic chip where *S. cerevisiae* or *S. pombe* cells grow in a single layer (adapted from [16]). When a cell is trapped below a PDMS pillar, all of its daughter cells are retained under the pillar and thus will form a colony (Fig. 1A, Supplementary Movie 1). After loading the microfluidic chip, we allowed the cells to adapt for at least three hours before we started time-lapse imaging in five minute intervals over a period of twelve hours. The average GR in each field of view can be calculated from all colonies where no cells reached the edge of the observation area during the duration of the experiment (Fig. 1B). The sensitivity of the assay is highest over a period of ~8 hours as only a few colonies outgrew the field of view. The obtained doubling time for *S. cerevisiae* is 123 ± 7 min and for *S. pombe* 128 ± 10 min (n = 5 different field of views, mean ± standard deviation), which is in agreement with commonly reported doubling times in liquid culture [17] (Supplementary Fig. 1 & 2). Our microscopy based assay is therefore suitable for measuring the GR.

**Figure 1:**
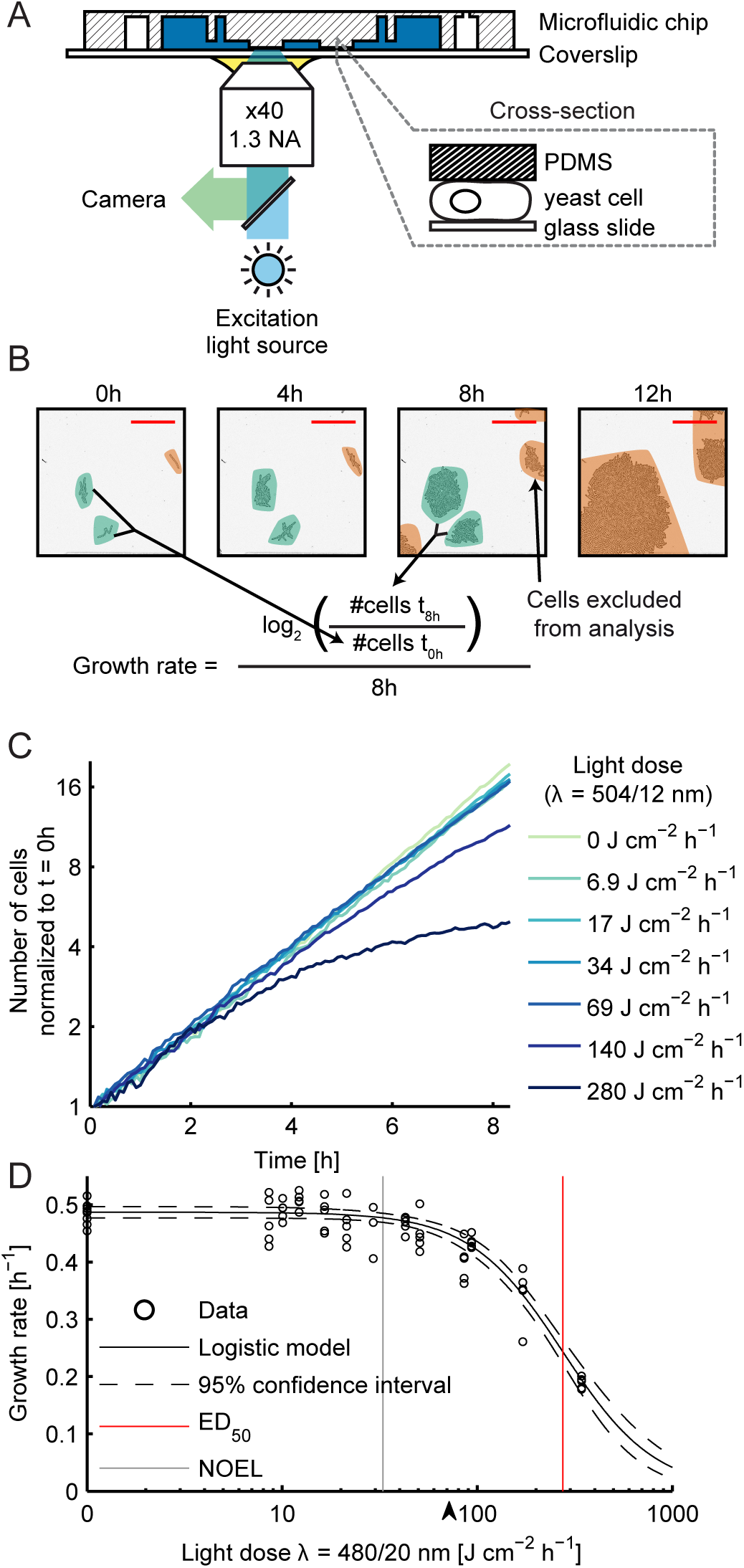
Measuring photomorbidity in a microfluidic chip using time-lapse imaging. A: Yeast cells are immobilized by clamping them between a PDMS pillar and a glass coverslip. B: Growth of *S. pombe* cells which are confined to grow in a single layer over 12 hours. The GR is calculated from the number of cells at the beginning and the end of an observation period. Colonies that reach the border of the field of view (orange regions) are excluded from the analysis. Red scale bars indicate 100 *μ*m. C: *S. cerevisiae* growth at seven different doses of teal excitation light. For each light dose, the GR was monitored on five independent culture pads. Cell numbers from each condition were summed and normalized to 1 for t=0 (n >93 for each condition). D: *S. cerevisiae* cells were exposed to different doses of cyan light. The GR obtained after 8 hours was plotted against the light dose (black circles) and the sigmoidal function (equation 2) was fitted to the data (black line). Dashed lines indicate the 95% confidence interval of the fit. The ED_50_ and the NOEL are indicated by red and grey lines, respectively. The black arrow indicates the lowest light dose at which stress could be detected using the general stress transcription factor Msn2 [12].

In a time-lapse microscopy experiment, the light dose per hour (*LD*, [J cm^−2^ h^−1^]) is the product of the exposure time (*t_E_*, [s]), the light intensity (*I*, [W cm^−2^]), and the imaging frequency (1/*t_Int_*, [h − 1], Supplementary Fig. 3):

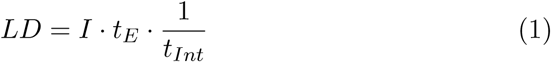

To accurately determine *LD*, we first measured the exact time a sample is exposed to light. This revealed that the sample is illuminated for an additional 187 ms compared to the set exposure time in our system (Supplementary Fig. 4), a hardware delay similar to previous reports [10]. Next, we determined the intensity of our light sources in combination with commonly used fluorescence imaging filter sets. For simplicity, we named the excitation wavelengths according to their perceived color. The main filters used are violet (λ = 390/18 nm), blue (438/24), cyan (480/20), teal (504/12), green (542/20) and red (600/14), and deviations are explicitly labeled (Supplementary Fig. 5, Supplementary Table 1 & 3). The measured intensities allowed us to determine the applied light doses of each color during our imaging experiments. To simplify the experimental setup, we used similar intensities (2.91 - 4.50 W cm^−2^) for all wavelengths in the photomorbidity assays except when indicated otherwise (Supplementary Table 5).

To test the effect of increasing doses of excitation light on the GR, we illuminated different culturing pads in the microfluidic device with different exposure times while keeping the light intensity and imaging interval constant (e.g. 7 light doses each repeated on 5 pads = 35 pads in one experiment). We found that the onset of growth retardation depended on the applied light dose in *S. cerevisiae*. At high doses, photomorbidity becomes visible within less than 3 hours while at lower doses, onset of photomorbidity can be delayed by several hours (Fig. 1C). We therefore calculated the average GR from the number of cells at the beginning (t=0 h) and end of the experiment (t=8 h). This allows for a sensitive and comparable measure of photomorbidity in cases of instantaneous and delayed growth retardation. The resulting dose-effect curve can be fitted using a sigmoidal function [18]:

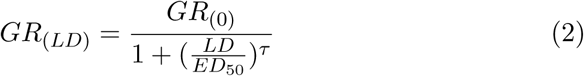

where ED_50_ is the effective dose at which the GR is reduced to 50% and τ is a measure of the capacity of cells to deal with increasing light doses (LD) once the GR is already reduced. For high values of τ, photomorbidity increased rapidly once a threshold dose was reached, whereas there was a slower increase in photomorbidity at small values of τ.

For practical purposes it is important to determine the highest light dose at which no adverse effects on the GR are observed, the so called no-observed effect level (NOEL). Small adverse effects seem to accumulate and are observable at later time points (Fig. 1C). We therefore fitted an exponential growth function to the first 3 hours of each pad of the teal assay and plotted the residuals over the whole time course (Supplementary Fig. 6). The resulting plot showed that once photomorbidity is induced, all pads behave similarly. Unfortunately, this way of determining the NOEL is impractical and we aimed to define the NOEL on a continuous scale. The intersection of the lower confidence bound of the dose effect curve and the non-illuminated control corresponds to a GR of 0.98. The resulting value indeed lies between the conditions where photomorbidity is observed in only a few pads and all pads and is therefore an experimentally useful definition. Determined in this way, the NOEL for cyan excitation light is 33 J cm^−2^ h^−1^ (Fig. 1D). This is lower than the lowest light dose (72 J cm^−2^ h^−1^) reported to increase nuclear localization frequency of Msn2 measured at a comparable wavelength (λ = 470/40 nm) [12] (black arrow in Fig. 1D), even though it is likely that the applied light doses in [12] were higher than reported due to a lack of determining hardware delays. Taken together, we operationally define that the NOEL corresponds to the light dose at which the fitted GR is reduced to 98% of the control.

### Morbidity is determined by the wavelength and light dose

We first asked whether the light doses that cause photomorbidity differ for different wavelengths and in different yeast species. We used the ED_50_ to evaluate the sensitivity of cells to the applied stress as this is a characteristic measure in toxicology [19]. *S. pombe* appears to be more sensitive to light than *S. cerevisiae* for all but blue excitation light (Fig. 2A). For both, *S. cerevisiae* and *S. pombe*, there is a general trend for longer wavelengths to cause less photomorbidity with the exception of teal excitation light (Supplementary Fig. 7 & 8). Photomorbidity was strongest in violet and blue light and least pronounced using green and red light. For red light applied to *S. cerevisiae*, we could not determine the ED_50_ as the low intensity of our light source lead to impractically long exposure times (Supplementary Table 1). It is important to note that the contribution of the brightfield illumination to the overall light dose can be neglected as it was ~50 fold lower than the lowest applied fluorescence excitation light dose throughout all experiments (brightfield dose = <18 mJ cm^−2^ h^−1^, lowest fluorescence excitation light dose = 890 mJ cm^−2^ h^−1^ violet). We conclude that each species has a specific sensitivity to light but the general trend is that longer wavelengths are better tolerated.

At the same cumulative dose we tested, whether changing intervals and intensities of the illumination light alters photomorbidity. We imaged *S. pombe* cells with teal light at intervals of 2.5, 5 or 7.5 minutes and found that the resulting ED_50_ values are not significantly different (Fig. 2B, Supplementary Fig. 9). Next, we illuminated *S. cerevisiae* cells with blue light of different intensities and observed that the ED_50_ values were not significantly different (Fig. 2C, Supplementary Fig. 10). We tested the effect varying excitation filter bandwidths on photomorbidity and illuminated *S. cerevisiae* cells with cyan light of similar central wavelength. Again, the resulting ED_50_ values were not significantly different (Fig. 2D, Supplementary Fig. 11). Previous studies indicated, that applying the excitation light using short pulses alters photomorbidity [20, 21]. We therefore illuminated *S. cerevisiae* with pulsed blue light (200kHz, 50% duty cycle) and again saw no change in the obtained ED_50_ (Supplementary Fig. 13). Our interpretation is that, at each excitation wavelength, the amount of light that induced a reduction in GR is solely determined by the cumulative light dose. This is in contrast to previous reports where high light intensities were shown to induce stronger adverse effects [9] [12]. However, this contradiction might be explained with hardware delays not taken into account in the aforementioned studies.

**Figure 2:**
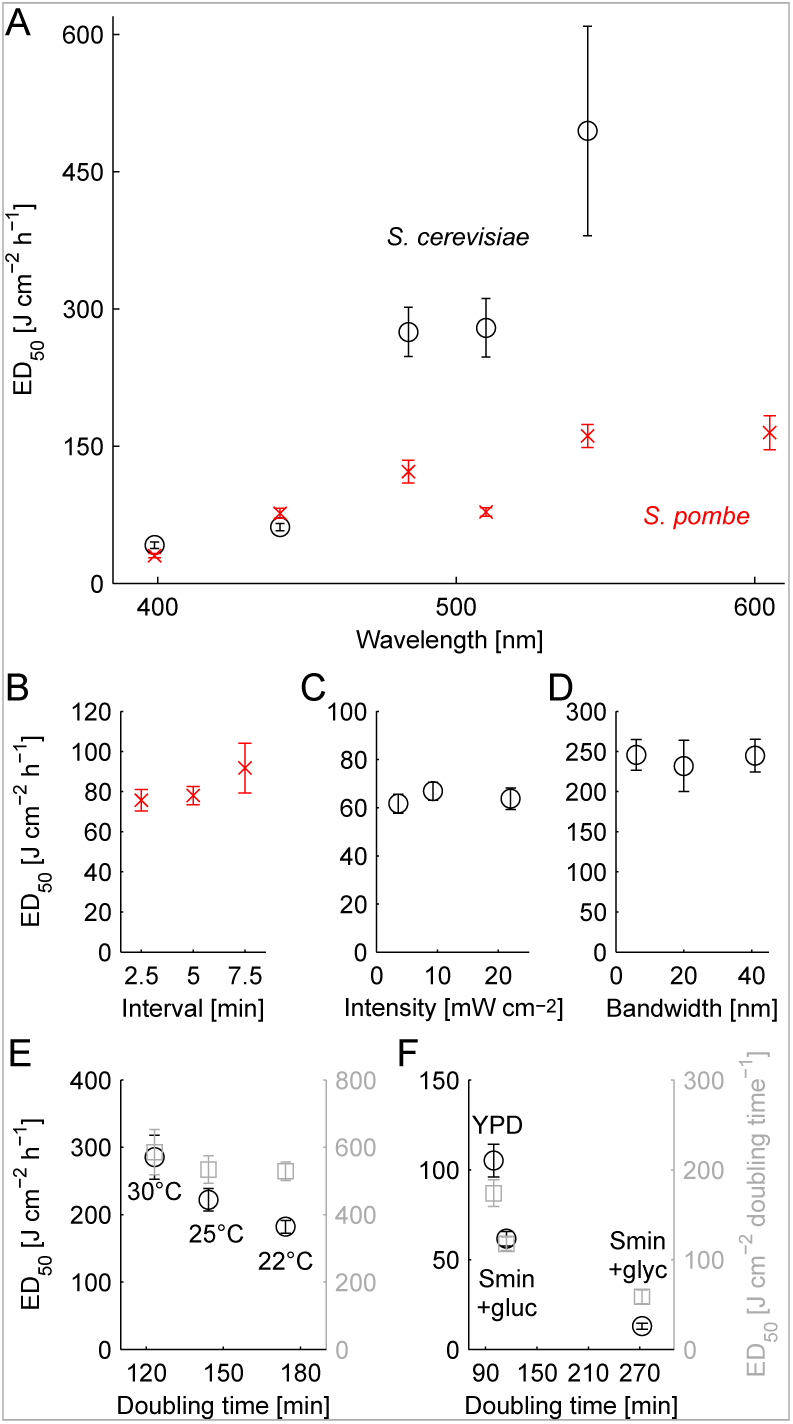
Sensitivity of *S. cerevisiae* and *S. pombe* to light using different imaging and culturing conditions. In each plot, the ED_50_-value with its 95% confidence interval are shown. A: Sensitivity of *S. cerevisiae* and *S. pombe* cells exposed to violet, blue, cyan, teal, green or red light (central wavelength, Supplementary Fig. 5). B: Sensitivity of *S. pombe* to teal excitation light at imaging intervals of 2.5, 5 or 7.5 minutes. C: Sensitivity of *S. cerevisiae* to blue light at light intensities of 3.8, 9.5 or 22 mW cm^−2^. D: Sensitivity of *S. cerevisiae* to cyan light using excitation filters with similar central wavelength but different bandwidths (488/6 nm, 480/20 nm, 494/41 nm). The light intensity was adjusted to 3.7 mW cm^−2^ for all three experiments. E: Sensitivity of *S. cerevisiae* to cyan excitation light at growth temperatures of 22, 25 and 30 °C. The ED_50_-values are plotted as light dose per hour (grey circles) and light dose per doubling time (black squares). F: Sensitivity of *S. cerevisiae* to blue excitation light in YPD, Smin+glucose or Smin+glycerol. ED_50_-values are plotted as in (E).

Photosensitivity of cells was shown to depend on their metabolic activity [22]. We first tested how slowed growth due to lower temperatures changes the susceptibility of cells to light. We grew *S. cerevisiae* at 30°C, 25°C or 22°C and illuminated them with cyan excitation light. The ED_50_ values were 290, 220 and 180 J cm^−2^ h^−1^, respectively (grey circles, Fig. 2E, Supplementary Fig. 14). However, when we normalized the applied light doses to the doubling time at the respective growth temperature, the ED_50_-values became 590 ± 67, 530 ± 40 and 530 ± 27 J cm^−2^ doubling time^−1^ and hence not significantly different from each other (black squares, Fig. 2E, Supplementary Fig. 14). Therefore, the light susceptibility at different temperatures can be extrapolated. This indicates that cells are able to tolerate a defined amount of light induced damage during the duration of a cell cycle.

If cells are able to tolerate a defined amount of light, then their susceptibility to light should depend on the nutritional growth conditions. To this end, we grew cells on rich media, minimal media with glucose and minimal media with the non-fermentable carbon source glycerol. Glycerol grown cells were twice as sensitive to blue light compared to glucose grown ones, even after normalization of the light dose to the doubling time (59 vs. 120 J cm^−2^ doubling time^−1^, black squares, Fig. 2F, Supplementary Fig. 15). Additionally, cells grown on glucose were more sensitive in minimal compared to rich media (120 vs. 170 J cm^−2^ doubling time^−1^, black squares, Fig. 2F, Supplementary Fig. 15). Taken together, changes in the metabolic flux lead to differences in photosensitivity which are not linearly related to the GR. These differences can be rationalized by changes in the expression of endogenous photosensitizers, like porphyrins, whose presence correlates with photosensitivity in *S. cerevisiae* [23] and which are upregulated in growth on non-fermentable carbon sources [24]. Depending on the absorption spectra of such photosensitizers the relative susceptibility at different wavelengths can change, as is the case for susceptibility to teal excitation light of cells grown in glycerol compared to glucose, which was even higher (170 vs. 550 J cm^−2^ doubling time^−1^, Supplementary Fig. 16).

Our results show that the sensitivity to growth inhibition by visible light depends on the light dose, wavelength, and metabolic state determined by the growth medium. For a given set of conditions, our results point to a tolerable light dose per cell cycle which is independent of the light intensity, imaging interval and bandwidth of the applied excitation light.

### Detectability of fluorescent proteins at NOEL light doses

The goal of live cell imaging is to obtain a detectable signal without damaging the cell. The obtained signal depends on the brightness of the fluorescent reporter and the amount of cellular autofluorescence [25]. We therefore judged the usability of different fluorescent proteins using the SNR [26]:

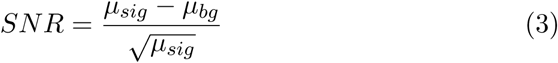

where *μ_sig_* is the *LD* dependent fluorescent signal and *μ_bg_* the textitLD dependent background fluorescence. 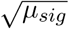 is an estimate of the statistical fluctuations in the measurement. The fluorescent signal is composed of the photons originating from the fluorescent probe (*P*) and the background fluorescence (*B*, cellular autofluorescence and fluorescence of media) as well as contributions from the noise of the camera 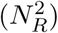 [25]. In short:

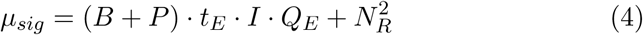

The magnitude of *P* and *B* depend on the quantity and brightness, and by how well the wavelength and bandwidth of the optical filters match the excitation and emission spectra of the fluorescent molecules. Additionally, *μ_sig_* depends on the quantum efficiency of the detector (*Q_E_*) in the associated spectral region and the exposure time *(t_E_*) and light intensity (*I*).

We first verified that the relationship between the obtained signal and the applied light dose is indeed linear and only depends on the concentration of the fluorophore. We imaged different yeast strains expressing Citrine as a C-terminal fusion to Vph1 (subunit of the vacuolar ATPase V0 domain), Cdc12 (a septin component) or Whi5 (a transcriptional repressor) with increasing exposure time (Fig. 3A). The calculated *in vivo* brightness (*μ_sig_* - *μ_bg_*) is 143 (Vph1), 43 (Cdc12) and 13 (Whi5) AU J^−1^ cm^2^ h. Next, we verified that the ratio between the *in vivo* brightness is independet of the used fluorophore by using mRuby2 and mKate2 as fluorescent tag. As long as *μ_sig_* is discernible from *μ_bg_*, the ratios are indeed constant (Supplementary Fig. 17, 18 & 19). In conclusion, each fluorescent protein has a specific *in vivo* brightness and can be compared between different fluorophores by using the background corrected slope.

The SNR is limited by “camera noise” at low light doses as *μ_sig_* is dominated by the readout noise of the camera 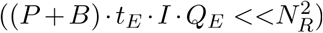. Once the signal is comparable to the noise, the SNR becomes limited by stochastic fluctuations in the fluorescent signal 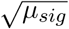 (”photon or shot noise”) [27]. We wanted to device a way by which we can predict the SNR based on a minimal set of measurements. We verified that the SNR can be estimated using the slopes of *μ_sig_* and *μ_bg_* (Equation 3) by comparing them to the SNR obtained from the single measurements (Fig. 3B). We denote them as eSNR (estimated, lines) and mSNR (measured, circles). Using the ratios described above, we tested whether the SNR can be predicted (pSNR). We tested the pSNR for Whi5 strains tagged with mRuby2 and mKate2 by using *μ*_whi5_ calculated from the previuously measured relative abundance (from Fig. 3A). The pSNR and mSNR values yield images with optically comparable quality (Fig. 3C). These results show that the light dose dependent image quality for a given fusion protein can be estimated, if *μ_sig_* and *μ_bg_* of a reference strain and the relative abundance of the fusion proteins are known.

**Figure 3:**
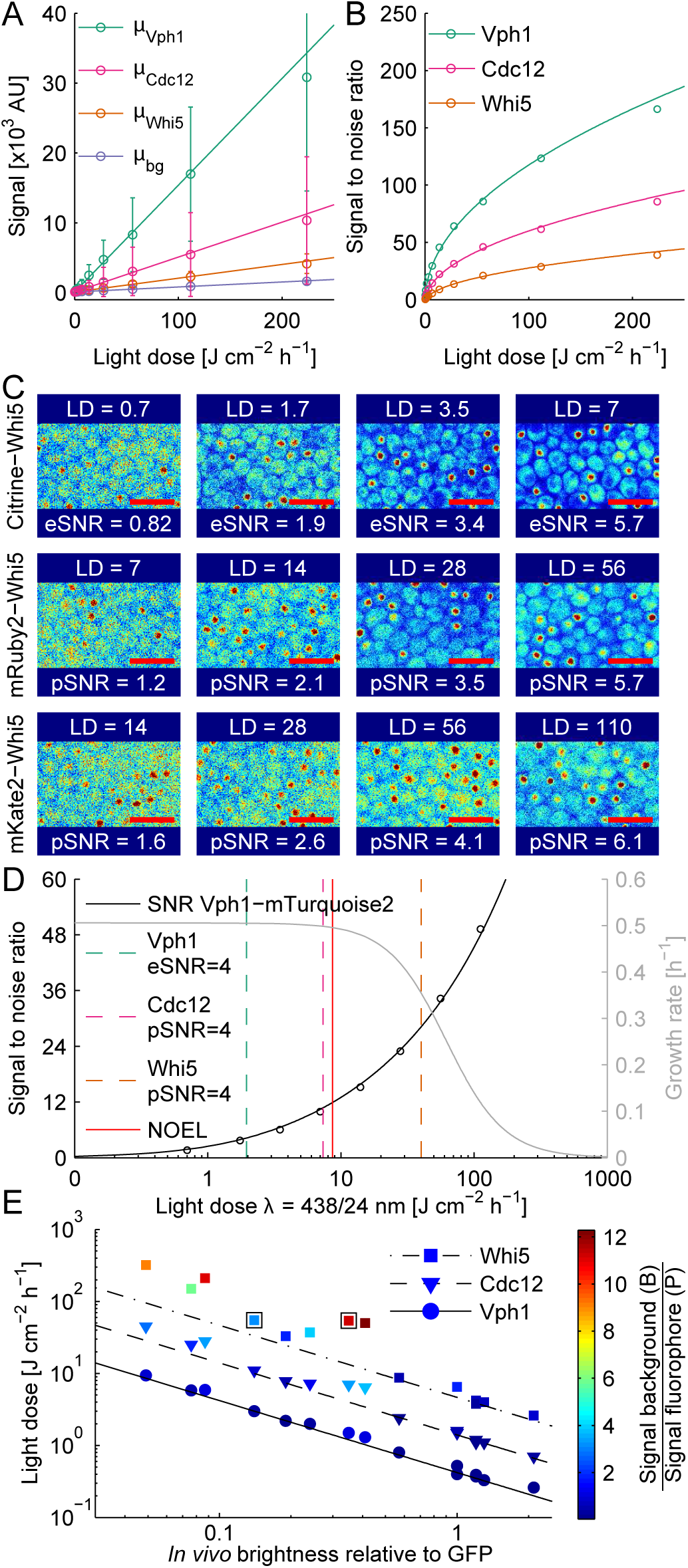
Relationship between the SNR, the light dose and photomorbidity. Detectability of fluorescent proteins at their respective NOEL in *S. cerevisiae* A: Signal intensity vs. excitation light dose for indicated strains tagged with Citrine. The mean ± SD of the cellular fluorescence is plotted. Points denote measurements and lines indicate linear fit. B: SNR vs. light dose plot. Points denote measured SNR values (mSNR) calculated using the single measurement points from A according to equation 3. Lines are the estimated SNR values (eSNR) calculated from the respective linear fit. C: Contrast adjusted false color fluorescent images of Whi5 strains tagged with different proteins at increasing light doses (LD in J cm^−2^ h^−1^). For Citrine the eSNRs is shown while the pSNR is shown for mRuby2 and mKate2 (see text for details). Contrast is adjusted such that 1% of the highest and lowest pixel values are saturated. Red scale bars indicate 10 *μ*m. D: SNR and photomorbidity vs. LD per hour of blue light with an imaging interval of 5 min for a Vph1-mTurquoise2 strain. Dashed green line indicates an eSNR of 4 for Vph1, while dashed red and orange lines indicate a pSNR of 4 for Cdc12 and Whi5, respectively. NOEL is indicated by a red line. E: LD required for an SNR of 4 vs. *in vivo* brightness for all tested fluorescent fusion proteins. The data points are colored according to the ratio of the background (untagged) divided by the fluorescent signal (tagged strain). The two boxes indicate the data points for Whi5-mKate2 and Whi5-mAmetrine.

The suitability of a fluorescent protein depends on its brightness and NOEL light dose. We decided to use an SNR of 4 at the NOEL light dose for comparison of commonly used fluorescent proteins (see Fig. 3C for image quality). We first measured the background corrected *in vivo* brightness of different fluorescent proteins for a set of Vph1-tagged strains (*μ_vph_*_1_). We then calculated the light doses required for SNR = 4 (Table 1) and extracted the expected GRs from the photomorbidity curves (Fig. 3D, Supplementary Fig. 20, Table 1). All tested Vph1 fusions could be detected at light doses below the NOEL in *S. cerevisiae*. In contrast, Whi5 could only be detected using fusion to GFP, mNeonGreen, Citrine, mRuby2, mKO*_κ_*, mKate2 and mCardinal fluorescent proteins without inducing photomorbidity. Additionally, mNeptune2 allows imaging of Cdc12 at NOEL light doses. As we could not determine a dose-effect curve for red excitation light, we utilized the data for green excitation light which leads to an overestimation of photomorbidity.

The dominant factor hindering high SNR imaging is the cellular autofluorescence (*B*). We noticed that Whi5-mKate2 and Whi5-mAmetrine yield similar SNRs at almost identical light doses (55 and 54 J cm^−2^ h^−1^), although mKate2 is only 40% as bright as mAmetrine. The main difference between the two imaging channels is the amount of cellular background fluorescence (*μ_bg_*, 43 and 4.8 AU J^−1^ cm^2^ h, Table 1). To better understand the interplay between fluorescent (*P*) and background signal (*B*), we plotted the light doses necessary to reach SNR = 4 against the *in vivo* brightness normalized to GFP (Fig. 3E). The two measures were inversely correlated for an abundant protein like Vph1. In contrast, the inverse correlation is lost when *B* is more than three times higher than *P* (colorbar in Fig. 3E). Knowing that the background fluorescence plays a major role in determining the SNR, we tried to find the best prediction for the amount of autofluorescence in a given imaging channel (Supplementary Fig. 23A). We found that the background fluorescence corrected for the bandwidth of the emission filter strongly correlated with the excitation wavelength (R^2^ = 0.92, Supplementary Fig. 23B). Therefore, the width of the emission filter can be used to optimize the ratio of *B* to *P*. Surprisingly, we found that the bandwidth corrected *B* is similar for channels excited at the same wavelength (compare Ex390/18 Em460/50 vs Em525/50 and Ex438/24 Em480/17 vs Em525/50, Supplementary Fig. 23C). However, as the background fluorescence of cells is influenced by the culturing conditions [28] and the imaging setup, its magnitude needs to be determined experimentally. In contrast to previous results, we did not see that a pulse-width modulated excitation light suppresses the background fluorescence 22).

**Table 1:** Properties of Vph1-fusion proteins. Light doses (LD) to reach SNR = 4 are given per hour based on an imaging interval of five minutes.

| Fluorescent protein | Excitation filter [nm] | Emission filter [nm] | Relative brightness | $\mu_{bg}$<br>[AU J<sup>-1</sup> cm<sup>2</sup> h] | LD<sub>vph1</sub> | LD<sub>cdc12</sub><br>at SNR=4<br>[J cm<sup>-2</sup> h<sup>-1</sup>] | LD<sub>whi5</sub> | GR<sub>vph1</sub> | GR<sub>cdc12</sub><br>at SNR=4<br>[%] | GR<sub>whi5</sub> | LD<sub>Bleach50</sub><br>[J cm<sup>-2</sup>] |
| --- | --- | --- | --- | --- | --- | --- | --- | --- | --- | --- | --- |
| tSapphire | 390/18 | 525/50 | 0.41 | 54 | 1.3 | 6.4 | 50 | 1.0 | 0.99 | 0.40 | 3010 |
| mAmetrine | 438/24 | 525/50 | 0.35 | 43 | 1.5 | 7.0 | 54 | 1.0 | 0.99 | 0.57 | 6320 |
| mTFP1 | 438/24 | 480/17 | 0.087 | 10 | 5.9 | 28 | 210 | 0.99 | 0.83 | 0.0079 | 866 |
| mTurquoise2 | 438/24 | 480/17 | 0.24 | 10 | 2.0 | 7.2 | 37 | 1.0 | 0.99 | 0.73 | 21200 |
| sfGFP | 488/6 | 525/50 | 1.0 | 20 | 0.4 | 1.6 | 6.5 | 1.0 | 1.0 | 1.0 | 5940 |
| eGFP | 488/6 | 525/50 | 1.0 | 20 | 0.52 | 1.5 | 6.5 | 1.0 | 1.0 | 1.0 | 15200 |
| mNeongreen | 488/6 | 525/50 | 2.1 | 20 | 0.26 | 0.7 | 2.6 | 1.0 | 1.0 | 1.0 | 6520 |
| mNeongreen | 504/12 | 542/22 | 1.2 | 8.8 | 0.37 | 1.2 | 4.2 | 1.0 | 1.0 | 1.0 | 3730 |
| CitrineA206K | 504/12 | 542/22 | 1.3 | 8.8 | 0.33 | 1.1 | 4.0 | 1.0 | 1.0 | 1.0 | 633 |
| mRuby2 | 561/4 | 600/32 | 0.19 | 3.8 | 2.2 | 7.8 | 33 | 1.0 | 1.0 | 1.0 | 5320 |
| tdKOkappa | 546/10 | 577/25 | 1.2 | 3.1 | 0.39 | 1.1 | 3.8 | 1.0 | 1.0 | 1.0 | 4200 |
| mKOκ | 546/10 | 577/25 | 0.57 | 3.1 | 0.80 | 2.4 | 8.7 | 1.0 | 1.0 | 1.0 | 7430 |
| mKate2 | 600/14 | 655/40 | 0.14 | 4.8 | 3.0 | 11 | 55 | 1.0 | 1.0 | 1.0 | 2930 |
| mCardinal | 600/14 | 655/40 | 0.076 | 4.8 | 5.8 | 25 | 150 | 1.0 | 1.0 | 1.0 | 7320 |
| mNeptune2 | 600/14 | 655/40 | 0.049 | 4.8 | 9.4 | 45 | 320 | 1.0 | 1.0 | >0.89 | 9100 |

### Predicting NOEL conditions for multi-color imaging

Currently, the extent to which cell physiology is affected by exposure to two or more separate excitation wavelengths is unknown. We first addressed whether photomorbid effects act independent of each other by exposing *S. cerevisiae* to combinations of two different excitation wavelengths. To score their interaction, we used the coefficient of drug interaction (CDI) which is a common pharmacological measure for the interaction of two treatments. The CDI is defined as:

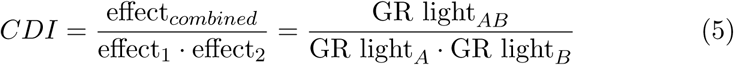

Treatments exhibiting additive behaviour are considered independent (CDIs ≈ 1), whereas synergistic or antagonistic behaviours yield CDIs of <0.8 or >1.2, respectively [29]. For example, we observed a GR reduction to 0.74 when combining blue and teal light. This corresponds to a CDI of 1 as the growth reduction of the single color treatment were 0.85 and 0.87, respectively. In general, photomorbidity seems to be additive since all tested pairwise combinations resulted in CDIs between 0.79 and 1.1 (Fig.4A, Supplementary Fig. 24). We confirmed this observation by using pairwise combinations of weakly morbid light doses in the more photosensitive yeast *S. pombe*. This resulted in CDIs between 0.91 and 1.1 (Fig. 4B, Supplementary Fig. 25). These results show that doses of excitation light relevant for non-toxic imaging act independently.

To achieve NOEL multi-color imaging, the combined effects of all used wavelengths should not reduce GRs to less than 0.98 of the control. Additivity implies that the multi-color NOEL for each individual wavelength can be calculated as the *n*th root of 0.98, where *n* denotes the number of distinct excitation wavelengths. We tested for the absence of photomorbid effects in three-color imaging (LDs for each excitation wavelength allow 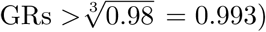 by applying combinations of excitation light doses aimed to detect a set of low abundant proteins in *S. cerevisiae*. First, we used a three color setup with negligible emission cross-talk (Citrine, mKO*κ* and mKate2) and simulated detection of two proteins expressed at the Whi5-level and one protein at the Cdc12-level each at an SNR of four. This combination resulted in GRs indistinguishable from the control (Fig. 4C). Next, we tested if we were accurately predicting the maximum available light dose for each single channel by adding a fourth light dose (mTurqouise with a relative GR of 0.99) and thereby slightly exceeding the predicted four-color NOEL. As expected, the resulting GR was reduced to 0.91 of the control. Taken together, our results show that the excitation light doses for unstressed multi-color imaging can be predicted from the single color photomorbidity curves.

**Figure 4:**
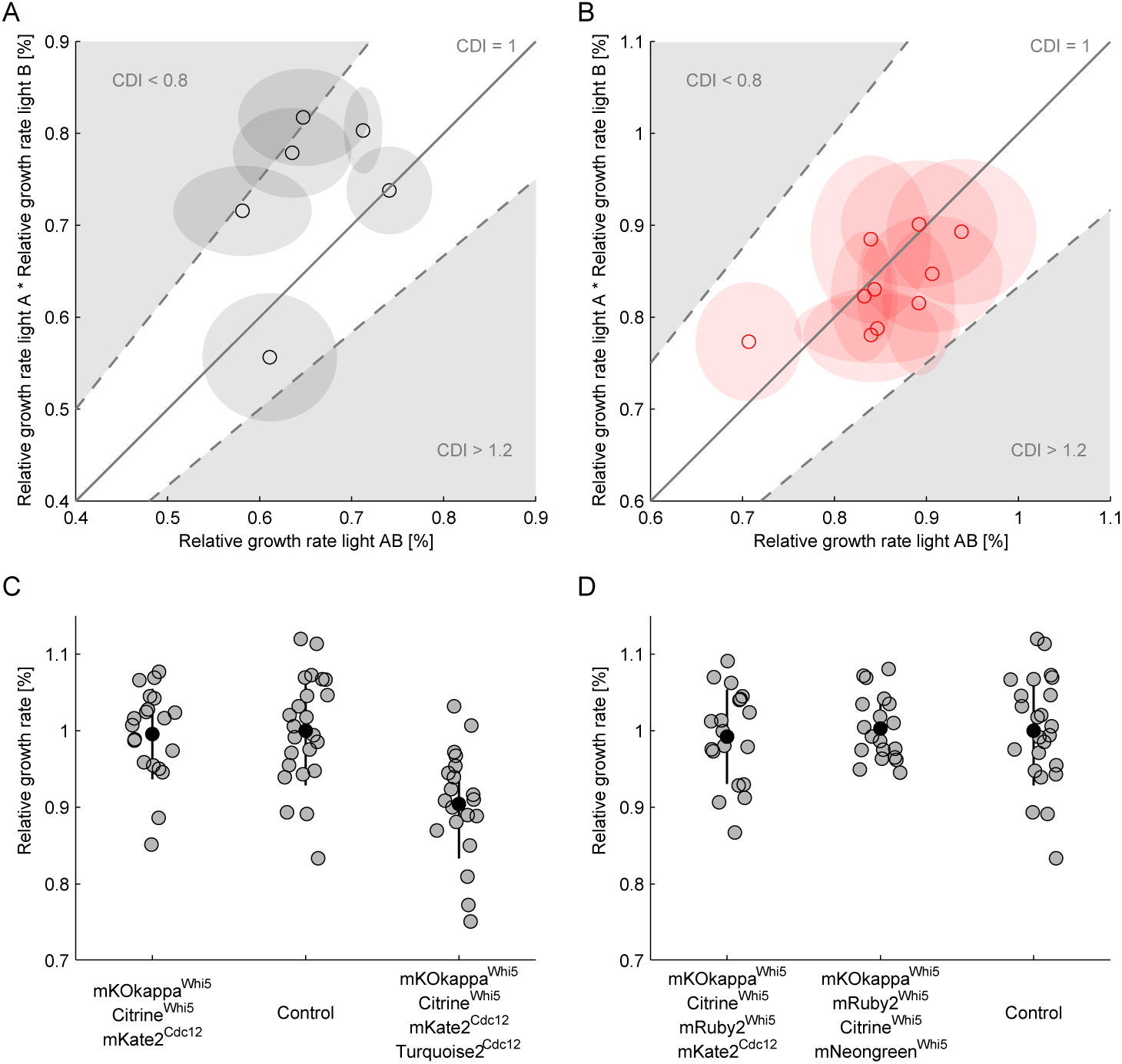
Photomorbidity is additive. A: Normalized *S. cerevisiae* GRs obtained during dual-color illumination with pairwise combinations of morbid light doses (GR light*_AB_*) were plotted against the product of the respective single color treatments (GR light*_A_* · GR light*_B_*). Each point is the mean of five pads and shaded circles denote standard deviations. For clarity, the line shows an additive model (CDI = 1) and the grey area where the CDI is either <0.8 or >1.2 (complete dataset in Supplementary Fig. 24). B: Same as in (A) but with *S. pombe* (complete dataset in Supplementary Fig. 25). C: NOEL imaging under the assumption of additive toxicity using four distinct excitation wavelengths. The mean normalized GR (black dots) from 20 pads (grey dots) and standard deviation (black lines) are plotted against the combination of the utilized fluorescence channels. The indication shows which target protein can be detected at an SNR of 4. D: NOEL imaging of 4 low expressed proteins when allowing for overlapping emission spectra. Axes as in (C).

To allow NOEL imaging of four low abundant proteins, it is conceivable to limit photomorbidity by exclusively using long excitation wavelengths. In this case, one has to allow for crosstalk of the respective emission spectra as well as using excitation wavelength bands which lie within the same photomorbidity window. We simulated an imaging combination (mKO*κ*, mRuby2, mKate2 and Citrine) which uses three distinct excitation bands (mKO*κ* and mRuby2 were excited within the green photomorbidity window). This would allow NOEL imaging of three proteins at the Whi5-level and one at the Cdc12-level (Fig. 4D). We next tested if a similar strategy was able to detect four low abundant proteins (Whi5-level). This is feasible using pairs of bright proteins excited with the same excitation wavelength band and substantial spectral overlap (Fig. 4D). Our results indicate that imaging using unconventional combinations of well performing fluorescent proteins has the potential to detect more low abundant targets without inducing photomorbidity. However, such experiments would require the use of spectral unmixing algorithms, due to the inherent emission crosstalk of the available fluorophores [30].

Since photomorbid effects of several wavelengths act additively, NOEL light doses for multi-color imaging are different than those for single-color imaging. The multi-color NOEL doses for each excitation wavelength can be calculated based on the respective photomorbidity curve. Together with the predicted light dose to reach a defined SNR, it is possible to determine a combination of fluorescent proteins that will allow unstressed imaging. Use of fluorescent reporters excited with short excitation wavelengths (<480 nm) should be limited to strongly expressed proteins or avoided completely whenever possible. Alternatively, wavelengths which cells are able to tolerate at large doses can be used for the detection of fluorescent proteins with overlapping excitation spectra.

## Discussion

To study the confounding effects of visible light in multi-color long term imaging, we first defined and characterized a sensitive, label-free and robust measure based on slowed cell growth (photomorbidity). We propose to use the term “photomorbidity”, defined as the decrease in growth rate due to light exposure, to describe sublethal effects of excitation light on dividing cells. This is consistent with the definition of “morbidity” by IUPAC: “departure, subjective or objective, from a state of physiological or psychological well-being” [19] and distinct from the term “phototoxicity”, which is used to describe cell death. We found that photomorbidity is specific for each species, wavelength, and metabolic condition and its onset can be delayed by several hours. We further found that, under a given set of conditions, cells appear to tolerate a specific amount of light per cell cycle, which is independent of the growth temperature, imaging frequency, light intensity and excitation filter bandwidth. Importantly, we found that the photomorbid effect of up to four excitation wavelengths is additive at light doses required for long-term microscopy experiments. Confounding effects in multi-color imaging can be avoided when combining light doses below the multi-color NOEL. We expect that these rules and our approach to establish unstressed imaging conditions can be applied to any dividing specimen and other imaging technologies. A detailed discussion of the practical aspects can be found in the supplementary material.

The choice of fluorescent proteins governs if a target protein can be detected with the required SNR at doses below the NOEL. This choice is based on the brightness of the fluorescent protein, the autofluorescence in the respective imaging channel and the photomorbidity exerted by the excitation light. Of course, the suitability of a fluorescent protein additionally depends on properties like oligomerization tendency, pH stability or maturation time [31]. However, we demonstrate that the photostability can be neglected in long-term imaging as the light doses leading to significant bleaching are above the NOEL (Table 1). The detectability of a fluorescent reporter can be optimized by choosing emission filters such that the contribution of the signal is maximized compared to the contribution of the background. Narrow excitation filters at the excitation maximum of the fluorophore can be used to optimize the excitation efficiency. Bright and well behaved fluorescent proteins with different stokes shifts emitting in the orange and red part of the spectrum would further facilitate unstressed multi-color imaging by replacing fluorescent reporters excited at the strongly photomorbid short wavelengths.

Our results challenge previously held beliefs in the field of fluorescence time-lapse imaging and allow for improvement of critical steps in established assay. Optimizing fluorescent filter sets based on our results is probably the most convenient and cost effective way to improve live-cell imaging experiments. We suggest to use excitation and emission filters with narrower bandwidth than commonly used. Additionally, our results highlight that extra care has to be taken to ensure NOEL imaging conditions for assays requiring UV-A excitation light or high magnification (Supplementary Fig. 21). Some of our results contradict previous studies, especially the absence of a correlation between photomorbidity and the intensity of the excitation light [9,12] and a lack of effect on photomorbidity and background fluorescence from pulsed illumination light [20,21]. The previously shown increase in phototoxicity at higher light intensities may be explained by unaccounted for hardware delays, which are easily overlooked but can add significantly to the real exposure time. In contrast, our inability to reproduce the effects of pulsed illumination shown in earlier reports cannot be easily explained and requires further investigation. Last, our experimental approach can be applied to any dividing specimen.

## Author contributions

All authors developed the concept and designed experiments. G.W.S and A.P.C performed phototoxicity experiments. All authors analyzed the phototoxicity data. G.W.S generated and characterized the fluorescent strains and analyzed the data. G.W.S. and F.R. wrote the manuscript. All authors read and approved the final manuscript.

## Acknowledgments

We thank Jörg Stelling for his support. We acknowledge Moritz Lang for the implementation of microfluidic pump control in the microscope software YouScope. We would like to thank Tina Sing, Roger Brent, Tania Roberts and Sven Rudolphi (Biochemical Research and Yeast End-Product Society) for discussions and critical reading of the manuscript.

## References

[1] Icha, J., Weber, M., Waters, J. C. & Norden, C. Phototoxicity in live fluorescence microscopy, and how to avoid it. Bioessays 39, 1–15 (2017).

[2] Laissue, P. P., Alghamdi, R. A., Tomancak, P., Reynaud, E. G. & Shroff, H. Assessing phototoxicity in live fluorescence imaging. Nat. Methods 14, 657–661 (2017).

[3] Cadet, J., Mouret, S., Ravanat, J. L. & Douki, T. Photoinduced damage to cellular DNA: direct and photosensitized reactions. Photochem. Photobiol. 88, 1048–1065 (2012).

[4] Sancar, A. Mechanisms of DNA repair by photolyase and excision nuclease. Angew. Chem. Int. Ed. 55, 8502–8527 (2016).

[5] Frigault, M. M., Lacoste, J., Swift, J. L. & Brown, C. M. Live-cell microscopy - tips and tools. J. Cell Sci. 122, 753–767 (2009).

[6] Kielbassa, C., Roza, L. & Epe, B. Wavelength dependence of oxidative DNA damage induced by UV and visible light. Carcinogenesis 18, 811–816 (1997).

[7] Godley, B. F. et al. Blue light induces mitochondrial DNA damage and free radical production in epithelial cells. J. Biol. Chem. 280,21061–21066 (2005).

[8] Tinevez, J.-Y. et al. A quantitative method for measuring phototoxicity of a live cell imaging microscope. Meth. Enzymol. 506, 291–309 (2012).

[9] Dixit, R. & Cyr, R. Cell damage and reactive oxygen species production induced by fluorescence microscopy: effect on mitosis and guidelines for non-invasive fluorescence microscopy. Plant J. 36, 280–290 (2003).

[10] Magidson, V. & Khodjakov, A. Circumventing photodamage in live-cell microscopy. Methods Cell Biol. 114, 545–560 (2013).

[11] Jacquet, M., Renault, G., Lallet, S., Mey, J. D. & Goldbeter, A. Oscillatory nucleocytoplasmic shuttling of the general stress response transcriptional activators Msn2 and Msn4 in Saccharomyces cerevisiae. J. Cell Biol. 161, 497–505 (2003).

[12] Logg, K., Bodvard, K., Blomberg, A. & Käll, M. Investigations on light-induced stress in fluorescence microscopy using nuclear localization of the transcription factor Msn2p as a reporter. FEMS Yeast Res. 9, 875–884 (2009).

[13] Lee, A. Y. et al. Mapping the cellular response to small molecules using chemogenomic fitness signatures. Science 344, 208–211 (2014).

[14] Dixon, S. J., Costanzo, M., Baryshnikova, A., Andrews, B. & Boone, C. Systematic mapping of genetic interaction networks. Annu. Rev. Genet. 43, 601–625 (2009).

[15] Carlton, P. M. et al. Fast live simultaneous multiwavelength four-dimensional optical microscopy. PNAS 107, 16016–16022 (2010).

[16] Frey, O., Rudolf, F., Schmidt, G. W. & Hierlemann, A. Versatile, simple-to-use microfluidic cell-culturing chip for long-term, high-resolution, time-lapse imaging. Anal. Chem. 87, 4144–4151 (2015).

[17] Sherman, F. Getting started with yeast. Meth. Enzymol. 350, 3–41 (2002).

[18] Hafner, M., Niepel, M., Chung, M. & Sorger, P. K. Growth rate inhibition metrics correct for confounders in measuring sensitivity to cancer drugs. Nat. Methods 13, 521–527 (2016).

[19] IUPAC glossary of terms used in toxicology - terms starting with M. https://sis.nlm.nih.gov/enviro/iupacglossary/glossarym.html (2016).

[20] Nishigaki, T., Wood, C. D., Shiba, K., Baba, S. A. & Darszon, A. Stroboscopic illumination usine light-emitting diodes reduces phototoxicity in fluorescence cell imaging. BioTechniques 41, 191–197 (2006).

[21] Boudreau, C. et al. Excitation light dose engineering to reduce photo-bleaching and photo-toxicity. Sci. Rep. 6, DOI: 10.1038/srep30892 (2016).

[22] Kvam, E. & Tyrrell, R. M. Induction of oxidative DNA base damage in human skin cells by UV and near visible radiation. Carcinogenesis 18, 2379–2384 (1997).

[23] Strakhovskaya, M. G., Shumarina, A. O., Fraikin, G. Y. & Rubin, A. B. Endogenous porphyrin accumulation and photosensitization in the yeast Saccharomyces cerevisiae in the presence of 2,2’-dipyridyl. J. Photochem. Photobiol. B, Biol. 49, 18–22 (1999).

[24] Roberts, G. G. & Hudson, A. P. Transcriptome profiling of Saccharomyces cerevisiae during a transition from fermentative to glycerol-based respiratory growth reveals extensive metabolic and structural remodeling. Mol. Genet. Genomics 276, 170–186 (2006).

[25] Pang, Z., Laplante, N. E. & Filkins, R. J. Dark pixel intensity determination and its applications in normalizing different exposure time and autofluorescence removal. J. Microsc. 246, 1–10 (2012).

[26] Waters, J. C. Accuracy and precision in quantitative fluorescence microscopy. J. Cell Biol. 185, 1135–1148 (2009).

[27] Lambert, T. J. & Waters, J. C. Assessing camera performance for quantitative microscopy. Methods Cell Biol. 123, 35–53 (2014).

[28] Heppert, J. K. et al. Comparative assessment of fluorescent proteins for in vivo imaging in an animal model system. Mol. Biol. Cell 27, 3385–3394 (2016).

[29] Bijnsdorp, I. V., Giovannetti, E. & Peters, G. J. Analysis of drug interactions. Methods Mol. Biol. 731, 420–434 (2011).

[30] Zimmermann, T. Spectral imaging and linear unmixing in light microscopy. Adv. Biochem. Engin./Biotechnol. 95, 245–265 (2005).

[31] Cranfill, P. J. et al. Quantitative assessment of fluorescent proteins. Nat. Methods 13, 557–563 (2016).

